# Laboratory Analysis of an Outbreak of *Candida auris* in New York from 2016 to 2018-Impact and Lessons Learned

**DOI:** 10.1101/760090

**Authors:** YanChun Zhu, Brittany O’Brien, Lynn Leach, Alexandra Clark, Marian Bates, Eleanor Adams, Belinda Ostrowsky, Monica Quinn, Elizabeth Dufort, Karen Southwick, Richard Erazo, Valerie B. Haley, Coralie Bucher, Vishnu Chaturvedi, Ronald J. Limberger, Debra Blog, Emily Lutterloh, Sudha Chaturvedi

**Affiliations:** Mycology Laboratory, Wadsworth Center, New York State Department of Health. Albany, NY 12208; Healthcare Epidemiology & Infection Control Program, New York State Department of Health, New Rochelle, NY 10801; Division of Healthcare Quality Promotion (DHQP), Centers for Disease Control and Prevention (CDC), Atlanta, GA 30333; Bureau of Healthcare Associated Infections, New York State Department of Health, Albany, NY 12237; Division of Epidemiology, New York State Department of Health, Albany, NY 12237; Department of Epidemiology and Biostatistics, School of Public Health, University at Albany, Albany, NY 12201; Department of Biomedical Sciences, School of Public Health, University at Albany. Albany, NY 12201

## Abstract

*Candida auris* is a multidrug-resistant yeast which has emerged in healthcare facilities worldwide, however little is known about identification methods, patient colonization, spread, environmental survival, and drug resistance. Colonization on both biotic and abiotic surfaces, along with travel, appear to be the major factors for the spread of this pathogen across the globe. In this investigation, we present laboratory findings from an ongoing *C. auris* outbreak in NY from August 2016 through 2018. A total of 540 clinical isolates, 11,035 patient surveillance specimens, and 3,672 environmental surveillance samples were analyzed. Laboratory methods included matrix-assisted laser desorption/ionization time-of-flight mass spectrometry (MALDI-TOF MS) for yeast isolate identification, real-time PCR for rapid surveillance sample screening, culture on selective/non-selective media for recovery of *C. auris* and other yeasts from surveillance samples, antifungal susceptibility testing to determine the *C. auris* resistance profile, and Sanger sequencing of ribosomal genes for *C. auris* genotyping. Results included: a) identification and confirmation of *C. auris* in 413 clinical isolates and 931 patient surveillance isolates, as well as identification of 277 clinical cases and 350 colonized cases from 151 healthcare facilities including 59 hospitals, 92 nursing homes, 1 long-term acute care hospital (LTACH), and 2 hospices, b) successful utilization of an in-house developed *C. auris* real-time PCR assay for the rapid screening of patient and environmental surveillance samples, c) demonstration of relatively heavier colonization of *C. auris* in nares compared to the axilla/groin, and d) predominance of the South Asia Clade I with intrinsic resistance to fluconazole and elevated minimum inhibitory concentration (MIC) to voriconazole (81%), amphotericin B (61%), 5-FC (3%) and echinocandins (1%). These findings reflect greater regional prevalence and incidence of *C. auris* and the deployment of better detection tools in an unprecedented outbreak.

## INTRODUCTION

*Candida auris*, a multidrug-resistant pathogenic yeast has emerged in healthcare facilities across the globe. *Candida auris* was first described as a new species in 2009 from the ear discharge of a hospitalized patient in Japan (1). In the last decade, cases of *C. auris* have been reported from 18 countries on five continents including Asia, Africa, Europe, South America, and North America (2–12). Fungemia caused by *C. auris* is associated with a high mortality rate and therapeutic failure (13). Ecological niches of *C. auris* remain unknown. However, successful colonization of human body sites and survival and persistence on surfaces within healthcare environments may be contributing to the outbreak and prevalence of *C. auris* worldwide. Whole genome sequence analyses of clinical *C. auris* isolates indicated that the emergence of clonal populations in Asia, South Africa, and South America occurred independently and spread locally within each region (7, 14, 15).

The identification of *C. auris* has been challenging for most clinical laboratories because of reliance on biochemical-based identification systems such as VITEK 2 and API 20C AUX. Due to the lack of *C. auris* in the database, these systems have misidentified it as *C. haemulonii* (5, 7, 9, 11, 14, 16). Other less commonly used biochemical platforms also misidentified *C. auris* as *Rhodotorula glutinis, C. famata, C. sake*, and *Saccharomyces cerevisiae* (16–18). Accurate identification of *C. auris* requires matrix-assisted laser desorption/ionization time-of-flight mass spectrometry (MALDI-TOF MS) (19) or sequencing of the 18S ribosomal gene (1). These reliable techniques are not readily available in most clinical laboratories. In the United States, the Food and Drug Administration (FDA) only recently approved the use of *C. auris* on the MALDI-TOF MS platform such as Bruker Daltonik, Bremen, Germany on April 24, 2018 (http://www.cidrap.umn.edu/news-perspective/2018/04/fda-approves-rapid-diagnostic-test-candida-auris), and bioMérieux VITEK MS, Marcy L’Etoile, France on December 21, 2018 (https://www.rapidmicrobiology.com/news/new-fda-clearance-for-vitek-ms-expanded-id-for-challenging-pathogens).

Apart from the difficult identification of *C. auris*, antifungal resistance testing has also been challenging. Approximately 90% of *C. auris* isolates are resistant to fluconazole (14). Moreover, elevated minimum inhibitory concentrations (MIC) for voriconazole have been reported for 50% of isolates, and higher MICs for amphotericin B for 30% of isolates (15, 16, 20). Resistance to echinocandins although low, has also been reported (20).

Adams and colleagues described the first 51 clinical cases and 61 colonized cases in the outbreak affecting in New York healthcare facilities (16). As part of an ongoing response to this *C. auris* outbreak 540 clinical isolates and 11,035 patient and 3,762 environmental surveillance samples have been collected in New York from 2016 through 2018. Here we provide the laboratory analysis of these specimens which highlight the application of rapid molecular screening to outbreak control, the unique characteristics of *C. auris* isolates, the spectrum of antifungal resistance, and the prevalence of other pathogenic yeasts.

## Materials and Methods

### Sample collection and case definitions

Clinical yeast isolates suspected of *C. auris* received from various healthcare facilities in NY from August 2016 to December 2018 were part of this investigation. However, surveillance samples (patient and environmental) collected from August 2016 to October 2018 from patients and their environment in various *C. auris*-affected healthcare facilities in NY were the major focus of these studies. Clinical cases were defined as the identification of a first *C. auris* isolate recovered from a specimen obtained to diagnose or treat disease (16). Colonized cases were defined as the recovery of the first *C. auris* isolate recovered from a sample for surveillance purposes (16). The environmental samples processed in this study were classified as porous (e.g. linen, carpet, etc.) and non-porous (e.g. metal knob, phone, TV monitor, bed rail, etc.) based on surface texture. A small percentage of environmental objects (2.8%) did not clearly fall within one of those two categories (e.g. chair, sofa, etc.) and were excluded from this study.

### Candida identification

*Candida* isolates were speciated by matrix-assisted laser desorption/ionization time-of-flight mass spectrometry (MALDI-TOF MS; Bruker, Bremen, Germany) using both the manufacturer and in-house validated library databases. The in-house library database was enriched by adding spectra of several *C. auris* isolates from the current NY outbreak and from the Centers for Disease Control and Prevention-Antibiotic Resistance (CDC AR) bank (https://www.cdc.gov/drugresistance/resistance-bank/index.html).

### Colonization study

<u>Sample types</u>: Axilla, groin, and nares swabs were collected individually or as a composite swab (axilla/groin or nares/axilla/groin) using the BD ESwab™ Liquid Amies Collection and Transport System (Becton Dickinson, Franklin Lakes, NJ, USA) from patients in various healthcare facilities. Occasionally, rectal or other body site swabs or body fluids were also collected. For environmental sampling, various objects and surfaces in healthcare facilities were swabbed with 3M Sponge Sticks (3M Health Care, St. Paul, MN, USA), and after an area was sampled, the sponge was removed from the collection stick and placed in zip-top bag. All surveillance swabs, other body fluids, and environmental sponges were transported to the laboratory within 48 to 72 hours at ambient temperature.

### Culture of swabs and sponges

Each ESwab containing 1 ml modified liquid Amies medium was vortexed for 30 seconds, and 50 μl was inoculated onto non-selective (Sabouraud dextrose agar containing antibacterials; SDA-A) and selective (Sabouraud dulcitol agar containing antibacterials and 10% salt; SDulA-AS) media as described previously (16). Two hundred microliters of the ESwab liquid were also inoculated in selective broth media minus agar mediaminus agar (SDulB-AS). Sputum and bronchial aspirates were streaked on all the media as described above. Urine samples if received in large volume (10 to 20 ml), were concentrated by centrifugation at 4000 RPM for 5 min; supernatant was decanted leaving about 3 ml; 100 μl was inoculated onto both SDA-A and SDulA-AS plates, and 1 ml was inoculated in 5 ml of SDulB-AS broth.

Each environmental sponge sample was placed in a Whirl-Pak Homogenizer Blender Filter bag containing 45 mL of phosphate-buffered saline (PBS) with 0.02% Tween 80. The bags were gently mixed in the Stomacher 400 Circulator (Laboratory Supply Network, Inc., Atkinson, NH, USA) at 260 rpm for 1 min, and the suspension was transferred into a 50-ml conical tube and centrifuged at 4,000 rpm for 5 min; supernatant was decanted, leaving about 3 ml of liquid at the bottom of the tube. The 3 mL liquid was vortexed briefly and of these 1 ml was removed, centrifuged, washed and re-suspended in 50 μl of PBS-BSA for DNA extraction as described below for swabs. From remaining 2 ml of liquid, 100 μl each was inoculated on SDA-A and SDulA-AS plates, and 1 ml was inoculated in 5 ml of SDulB-AS broth.

Agar plates and broth tubes were incubated at 40°C for a maximum of two weeks.

### Enumeration of colony forming units (CFU)

To determine the extent of colonization on skin by *C. auris*, the colonies recovered on the plates were counted. If colonies were numerous, a 10-fold dilution series of the swab was prepared and plated for colony counts. Recovered colonies were identified by MALDI-TOF MS and results were expressed as colony forming units (CFU) per swab.

### Real-time PCR

A real-time PCR assay developed in the laboratory (21) for the rapid screening of surveillance samples for *C. auris*, was deployed effective May 17, 2017 following approval from the New York State Clinical Laboratory Evaluation Program. In brief, 200 μl of ESwab liquid Amies and 1 ml of concentrated liquid following sponge processing were washed twice with phosphate buffered saline (PBS) containing 0.1% bovine serum albumin (BSA), re-suspended in 50 μl of PBS-BSA followed by freezing, heating, bead-beating, and centrifugation at 13,000 RPM for 5 min, and 5 μl of the extracted DNA was tested in duplicate on the real-time PCR assay. A cycle threshold value of ≤37 was reported as positive, and >37 was reported as negative for *C. auris*. If PCR inhibition was observed, the results was reported as inconclusive (21).

### Antifungal susceptibility testing of C. auris and other closely related species

The minimum inhibitory concentrations (MICs) of azoles and echinocandins were determined by custom TREK frozen broth microdilution panels (catalog no. CML2FCAN; Thermo Fisher Scientific, Marietta, OH, USA), and MICs of amphotericin B and 5-flucytosine (5-FC) were determined by Etest as recommended by the manufacturer (AB Biodisk; bioMérieux, Solna, Sweden) except that MICs were read at 24 h post-incubation or until a confluent lawn of growth was seen (16). There are currently no established *C. auris*-specific susceptibility breakpoints. Therefore, CDC defined breakpoints were used for azoles, amphotericin B and echinocandins, and also MIC value of 1.5 for amphotericin B was rounded up to 2.0 (https://www.cdc.gov/fungal/candida-auris/c-auris-antifungal.html).

### Phylogenetic analysis of Candida auris

Genomic DNA of *C. auris* from the current outbreak and reference strains procured from CDC AR bank was extracted using a QIAcube DNA extractor with the QIAamp DNA Mini Kit (Qiagen, Hilden Germany). The extracted DNA was amplified for the internal transcribed spacer (ITS) and D1/D2 regions of the ribosomal gene with primer set ITS1-ITS4 and NL1-NL4, respectively [<u>44</u>]. Conventional PCR was performed using proof reading AccuTaq ™ LA DNA Polymerase (Sigma-Aldrich, St. Louis, MO, USA) with initial denaturation at 95°C for 3 min, followed by 30 cycles of denaturation at 94°C for 1 min, annealing at 55°C for 1 min, and extension at 68°C for 3 min and the final extension at 68°C for 10 min. The PCR amplicons were sequenced, assembled, and edited using Sequencher 5.0 software (Gene Codes Corp., Ann Arbor, MI, USA) and BLAST searched against two databases: GenBank (www.ncbi.nlm.nih.gov/) and CBS-KNAW (www.cbs.knaw.nl/). Multiple alignments of ITS and D1/D2 sequences of *C. auris* from the current outbreak and reference strains were done using the Geneious R9 version 9.1.6 (Biomatters, Inc., Newark, NJ) and the phylogenetic analysis of the aligned sequences was done using the neighbor-joining (NJ) method with 2,000 bootstrap replicates on the same software.

### Statistical analysis

GraphPad Prism 8 software for Mac (GraphPad Software, Inc., La Jolla, CA) was used for statistical analysis of the results. Two by two tables and the nonparametric Mann Whitney test were used to compare samples.

### Nucleotide Sequence Accession Numbers

All nucleotide sequences of *C. auris* were deposited in GenBank under accession numbers: ITS sequences: MN338097-MN338196; D1/D2 sequences: MN337432-MN337531.

### Results communication

Any positive real-time PCR result was immediately communicated to New York State Department of Health (NYSDOH) epidemiologists and the requesting facilities, allowing facilities to rapidly implement an infection control response. Following real-time PCR, culture results and then antifungal susceptibility testing results were communicated. Any unusual results such as resistance to echinocandins, were also communicated to the Centers for Diseases Control and Prevention (CDC) as an alert.

## RESULTS

### New York state-wide Candida auris identification

We received 540 isolates of yeasts suspected of being *C. auris* from various clinical, public, and commercial laboratories. Of these, 413 were confirmed as *C. auris*, 12 as *C. duobushaemulonii*, and 7 as *C. haemulonii* (Supplementary Table 1). The remaining 108 isolates were confirmed as other yeasts (data not shown). In most cases, there was no correlation seen between presumptive identification (ID) by the requesting laboratory and the method they used for identification. Laboratories using ITS sequencing provided a 100% correct identification of *C. auris* (42/42) while laboratories using Bruker MALDI provided a 72% (106/147) correct identification. Laboratories using the VITEK MS provided 21% (9/43) correct identification of *C. auris.* Interestingly, some clinical laboratories using the biochemical-based system, VITEK 2 provided a presumptive *C. auris* identification. Whether these laboratories presumed identification of *C. haemulonii* as *C. auris* or used a ‘research use only’ database of MALDI-TOF from Bruker or Biomerieux is unclear.

### Clinical cases

A total of 277 clinical cases were confirmed by culture during this period. The majority of *C. auris* isolates from clinical cases were recovered from blood (51%) followed by urine (23%) and then other body sites (Table 1). Based on these results, a blood stream infection (BSI) was by far the most common clinical infection. An additional 136 *C. auris* isolates were recovered from some of these 277 clinical case patients, resulting in a total recovery of 413 *C. auris* isolates.

**Table 1.**
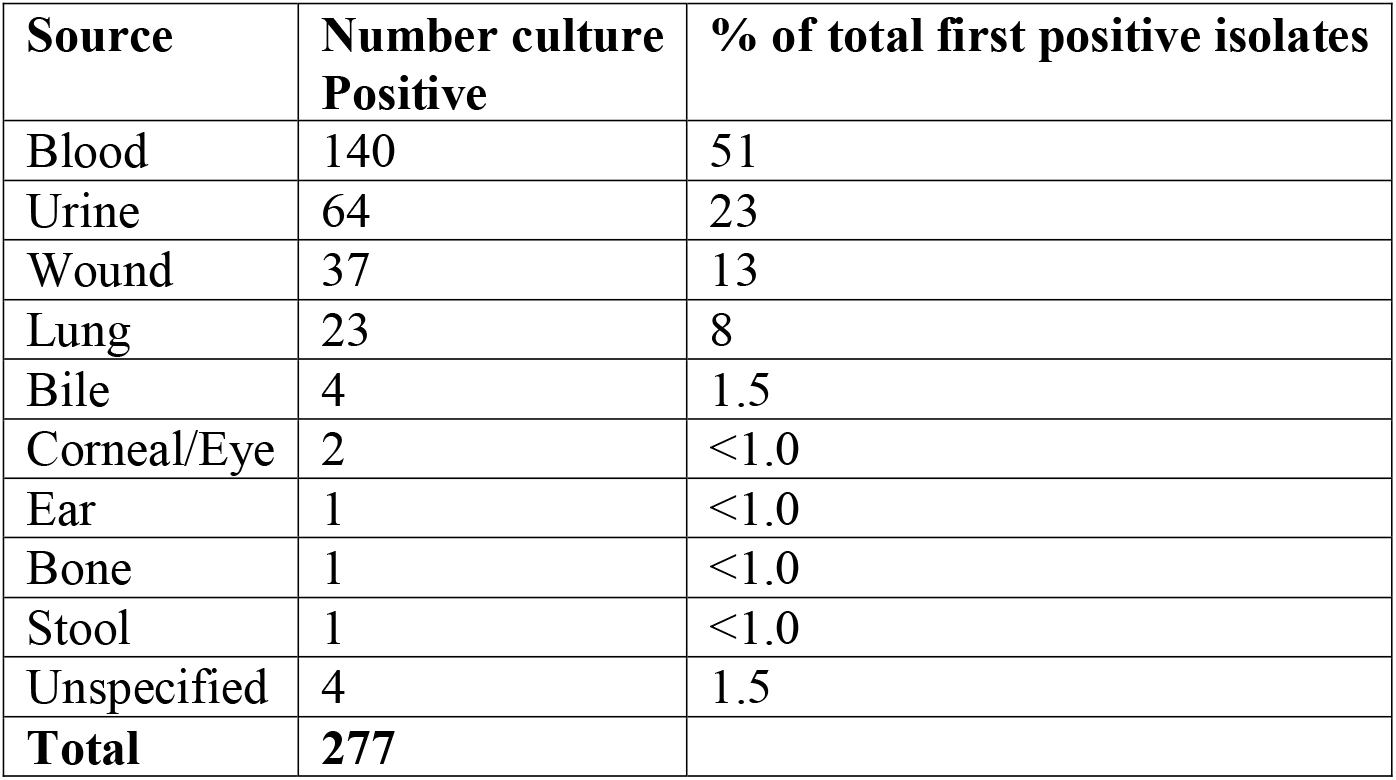
Source of first *Candida auris* isolate to define a clinical case.

| Source | Number culture Positive | % of total first positive isolates |
| --- | --- | --- |
| Blood | 140 | 51 |
| Urine | 64 | 23 |
| Wound | 37 | 13 |
| Lung | 23 | 8 |
| Bile | 4 | 1.5 |
| Corneal/Eye | 2 | <1.0 |
| Ear | 1 | <1.0 |
| Bone | 1 | <1.0 |
| Stool | 1 | <1.0 |
| Unspecified | 4 | 1.5 |
| **Total** | **277** |  |

### Colonized cases

Nine hundred thirty-one (8.4%) of 11,035 patient surveillance samples tested positive for *C. auris* by culture (Table 2). These included 450 first positive patient surveillance samples representing 350 colonized cases, 183 subsequent positive patient surveillance samples from some of the 350 colonized cases, and 298 positive patient surveillance samples from known clinical cases. A total of 151 healthcare facilities in which the patients received care in the 90 days prior to their *C. auris* diagnosis were affected by the outbreak, and these included 56 hospitals, 92 nursing homes, 1 long-term acute care hospital (LTACH) and 2 hospices.

**Table 2.** *C. auris* culture from surveillance samples tested from August 2016 to October 2018

| Source | Samples culture Positive/Samples Tested (%) |
| --- | --- |
| Axilla/Groin | 231/3928 (6.0) |
| Nares | 275/4058 (6.8) |
| Nares/Axilla/Groin | 121/1846 (6.6) |
| Axilla | 68/340 (20.0) |
| Groin | 70/311(22.5) |
| Rectal | 37/165 (22.4) |
| Wound | 69/187 (36.9) |
| Urine | 20/57 (35.1) |
| Respiratory | 18/51(35.3) |
| Ear | 6/15 (40.0) |
| Skin | 5/47 (8.8) |
| Unspecified | 11/30 (36.7) |
| **Total** | **931/11035 (8.4%)** |

Of 624 surveillance samples cultured, 450 were positive defining 350 colonized cases (Table 3). Although first positive surveillance sample defining 350 colonized cases originated from different body sites, we chose axilla/groin and nares to understand the probability and the extent of *C. auris* colonization as they made up the bulk of surveillance samples. Of 222 axilla/groin samples tested, 178 (80%) were positive for *C. auris* while of 215 nares samples tested, 125 (58%) were positive for *C. auris* (Table 3). When the extent of *C. auris* colonization in 178 axilla/groin and 125 nares positive sites were analyzed randomly; nares harbored 2 logs (P<0.001) higher *C. auris* than the axilla/groin (Fig. 1A). When 74 of axilla/groin and nares in parallel (from same patient) were analyzed; nares harbored 2 logs higher *C. auris* than the axilla/groin (Fig. 1B). These results indicated that axilla/groin is the preferred site of *C. auris* colonization as compared to nares in the colonized patients, but if nares is colonized, it harbors relatively higher burden of *C. auris* than axilla/groin. These results plus practical logistic and resource issues prompted us to use one composite swab of nares/axilla/groin for determining colonized cases, effective January 2018 for all point prevalence studies. Of 350 colonized cases, 106 were indeed identified using one composite swab of nares/axilla/groin in the present investigation.

**Table 3.** Source of first *C. auris* isolate to define a colonized case

| **Source** | **Samples**<br>**Positive/Samples**<br>**Tested** | **% of total**<br>**first positive**<br>**samples** |
| --- | --- | --- |
| Axilla/Groin (bilateral) | 178/222 | 80 |
| Nares (bilateral) | 125/215 | 58 |
| Nares/Axilla/Groin (bilateral) | 106/106 | 100 |
| Axilla (unilateral) | 10/20 | 50 |
| Groin (unilateral) | 10/20 | 50 |
| Nares (unilateral) | 6/14 | 43 |
| Wound | 4/11 | 36 |
| Rectal | 4/7 | 57 |
| Ear | 4/6 | 67 |
| Skin | 1/1 | 100 |
| Not specified | 2/2 | 100 |
| **Total** | **450/624** | **72%** |

**Fig. 1.**
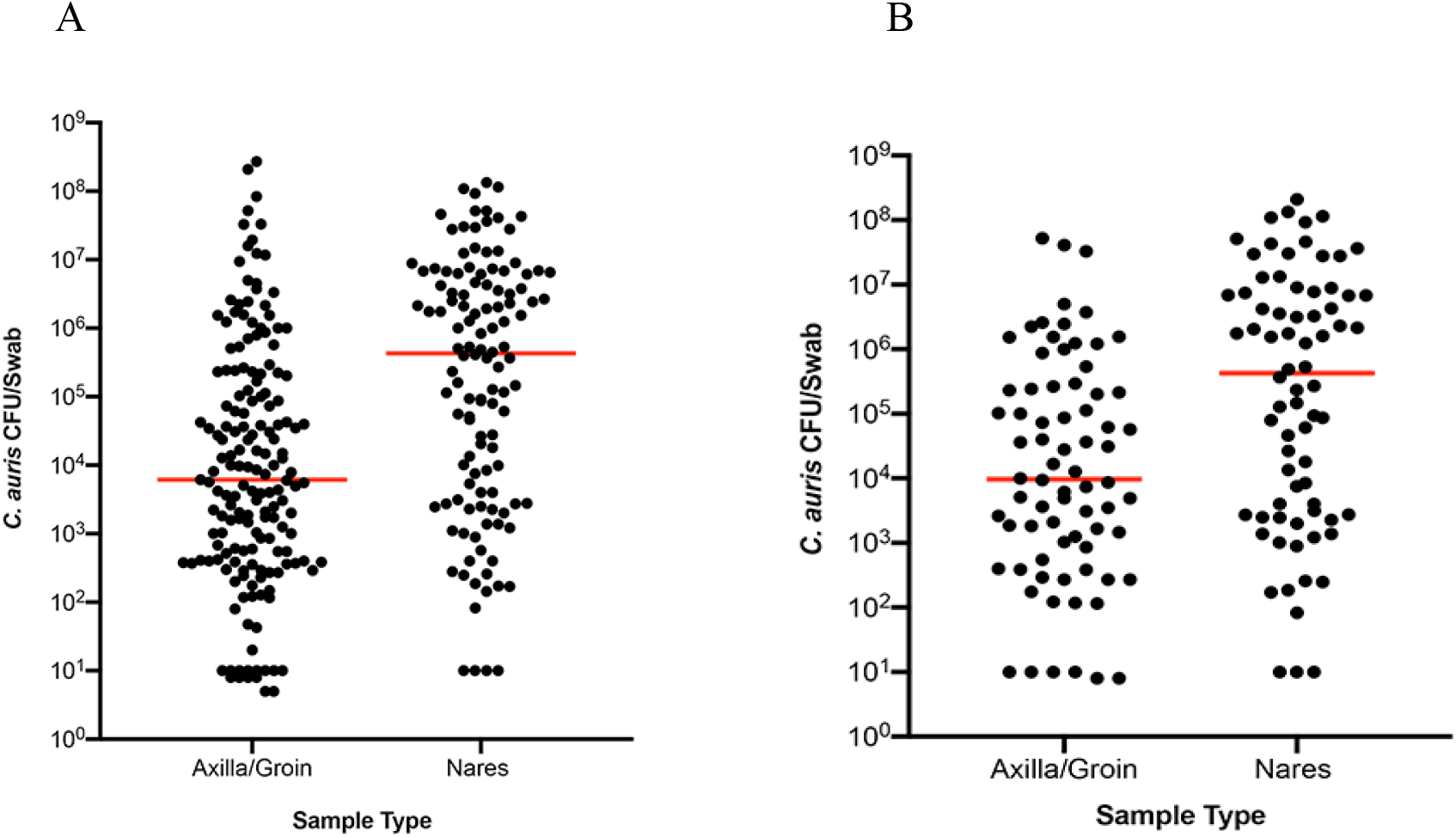
Colonization of the axilla/groin and nares. Axilla/groin and nares were swabbed, processed for culture, and recovered colonies were counted and results were expressed as CFU/swab. Each dot represents total CFU/swab/patient and the horizontal bar within each group represents the median. A. unpaired samples from patients colonized with *C. auris*. B. Paired samples from patients colonized with *C. auris*. In both cases, the median *C. auris* CFU was 2-logs higher in nares than in axilla/groin (p<0.001).

### Utility of C. auris real-time PCR assay in patient surveillance screening

Following NYSDOH CLEP approval of *C. auris* real-time PCR assay for patient surveillance samples effective May 2017, a total of 9,982 patient samples were tested from May 2017 to October 2018, and 6,834 of those samples were cultured as part of point prevalence studies. In comparison to culture as a gold standard, the diagnostic accuracy, sensitivity and specificity of the real-time PCR assay was 98.36%, 93.32% and 98.38%, respectively (Table 4 A). The performance was consistent with findings in our earlier investigation (21). ‘

**Table 4 A.**
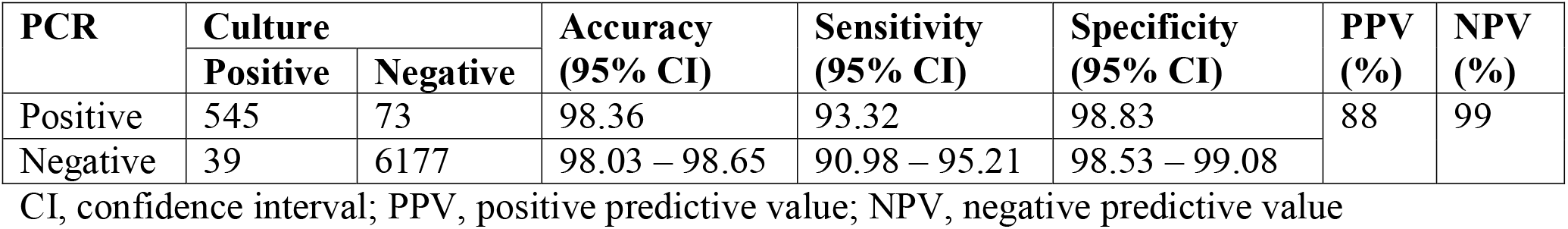
Efficacy of the *C. auris* real-time PCR assay for patient surveillance samples

### Utility of C. auris real-time PCR assay in environmental surveillance screening

Four hundred thirty-four of 3,672 environmental samples were positive by real-time PCR (11.8%). *Candida auris* was cultured from 109 of 434 PCR positive samples (25%). The diagnostic accuracy, sensitivity and specificity of the real-time PCR assay for the environmental samples was 69.42%, 98.63% and 67.5%, respectively (Table 4 B). *C. auris* colonization of culture positive environmental surfaces as quantified by CFU, found a range of concentrations ranging from a low of <50 CFU/surface to a high of >10^5^ CFU/surface; degree of colonization similar irrespective of porous or non-porous surface (Fig. 2).

**Table 4 B.** Efficacy of the *C. auris* real-time PCR assay for environmental surveillance samples

| PCR | Culture |  | Accuracy<br>(95% CI) | Sensitivity<br>(95% CI) | Specificity<br>(95% CI) | PPV<br>(%) | NPV<br>(%) |
| --- | --- | --- | --- | --- | --- | --- | --- |
|  | Positive | Negative |  |  |  |  |  |
| Positive | 72 | 362 | 69.42 | 98.63 | 67.5 | 16.67 | 99.87 |
| Negative | 1 | 752 | 66.72 - 72.09 | 92.60 - 99.97 | 64.67-70.25 |  |  |

CI, confidence interval; PPV, positive predictive value; NPV, negative predictive value

**Figure 2.**
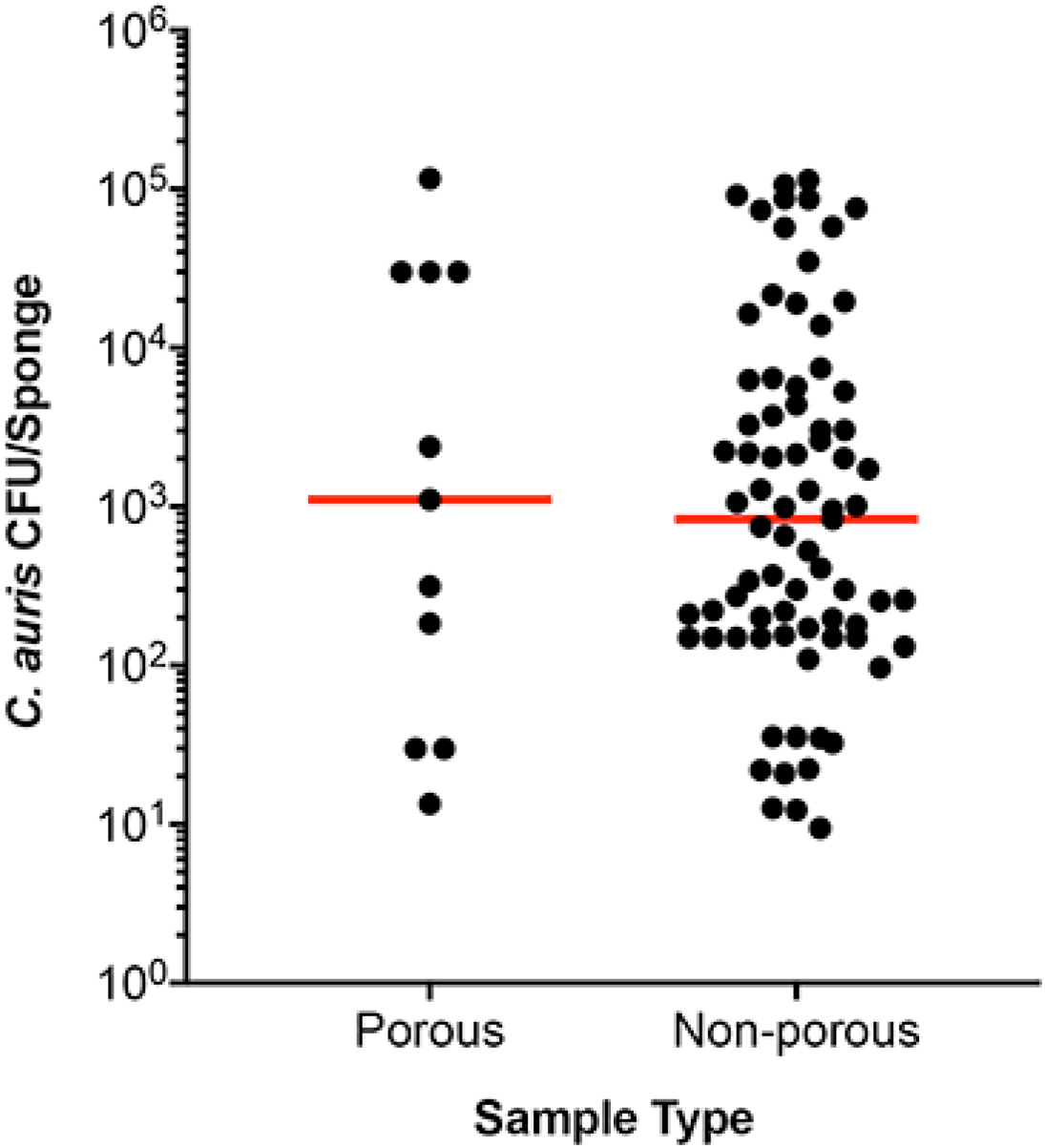
Colonization of environmental surfaces with *C. auris*. Environmental surfaces of various healthcare facilities were sponge swabbed and processed for culture, and recovered colonies were counted and results were expressed as total CFU/sponge. The environmental surfaces were divided into porous (i.e. linen, carpet) and non-porous (i.e. plastic & metal devices) for data analysis. Each dot represents total CFU recovered/per area swabbed, and the horizontal bar within each group represents the median. The median *C. auris* CFU was approximately 10^3^ irrespective of the surface type tested.

### Multiple alignment and phylogenetic analysis of ribosomal genes of C. auris

To determine the genetic makeup of *C. auris* causing the current NY outbreak, ITS and D1/D2 regions of the ribosomal gene were used. The ITS conventional PCR yielded a 400-bp amplicon and following sequencing and trimming of 5’ and 3’ prime ends of the ITS gene, a 311-bp sequence was used for the multiple alignment. There were four mutations at the 5’end of the gene at positions 67 (A to C), 68 (C to T), 70 (A to T), 74 (C to G), and three mutations at the 3’ prime end of the gene at position 307 (T to C), 308 (C to G), 309 (G to T) between South Asia Clade I and East Asia Clade II isolates. Additionally, two nucleotide insertions were found at positions 75 and 76 of A & T in South Asia clade I, which were absent in East Asia clade II. The phylogenetic analyses using a neighbor joining method revealed well separation of these two clades from each other (Fig. 3A).

**Figure 3 A.**
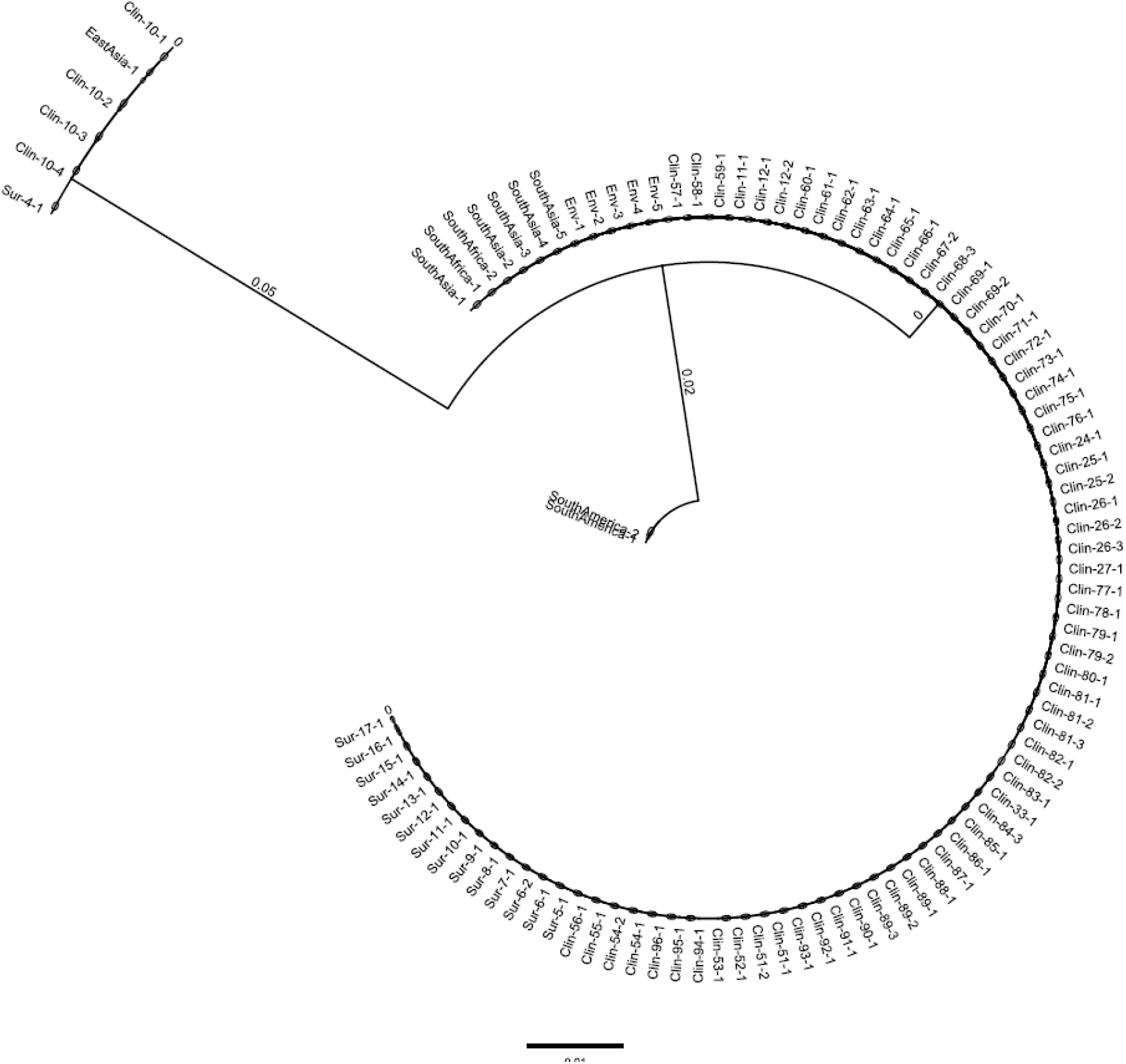
Phylogenetic analysis of *C. auris* isolates from NY using ITS sequences. ITS nucleotide sequences of 622 isolates of *C. auris* recovered from clinical patients (Clin), colonized patients (Sur), environmental surfaces (Env), and 10 standard isolates from the CDC-AR bank were aligned. The neighbor joining method was used to construct the phylogenetic tree. The bootstrap scores are based on 2,000 reiterations. The outbreak was predominately the South Asia Clade I with a minor population of the East-Asia Clade II. A tree representing a few *C. auris* isolates from each group is shown. Note, ITS could not distinguish South Africa Clade III (CDC AR) from South Asia Clade I.

Amplification of the D1/D2 region revealed a 500-bp amplicon and following sequencing and trimming of 5’ and 3’ primed ends, a 426-bp sequence was used for the multiple alignment. There were two mutations at position 372 and 389 of T to C, and an insertion at position 392 and 393 of two Ts in South Asia clade I versus East Asia clade II. The topology of the neighbor joining tree was similar to that of ITS (Fig. 3B).

**Figure 3 B.**
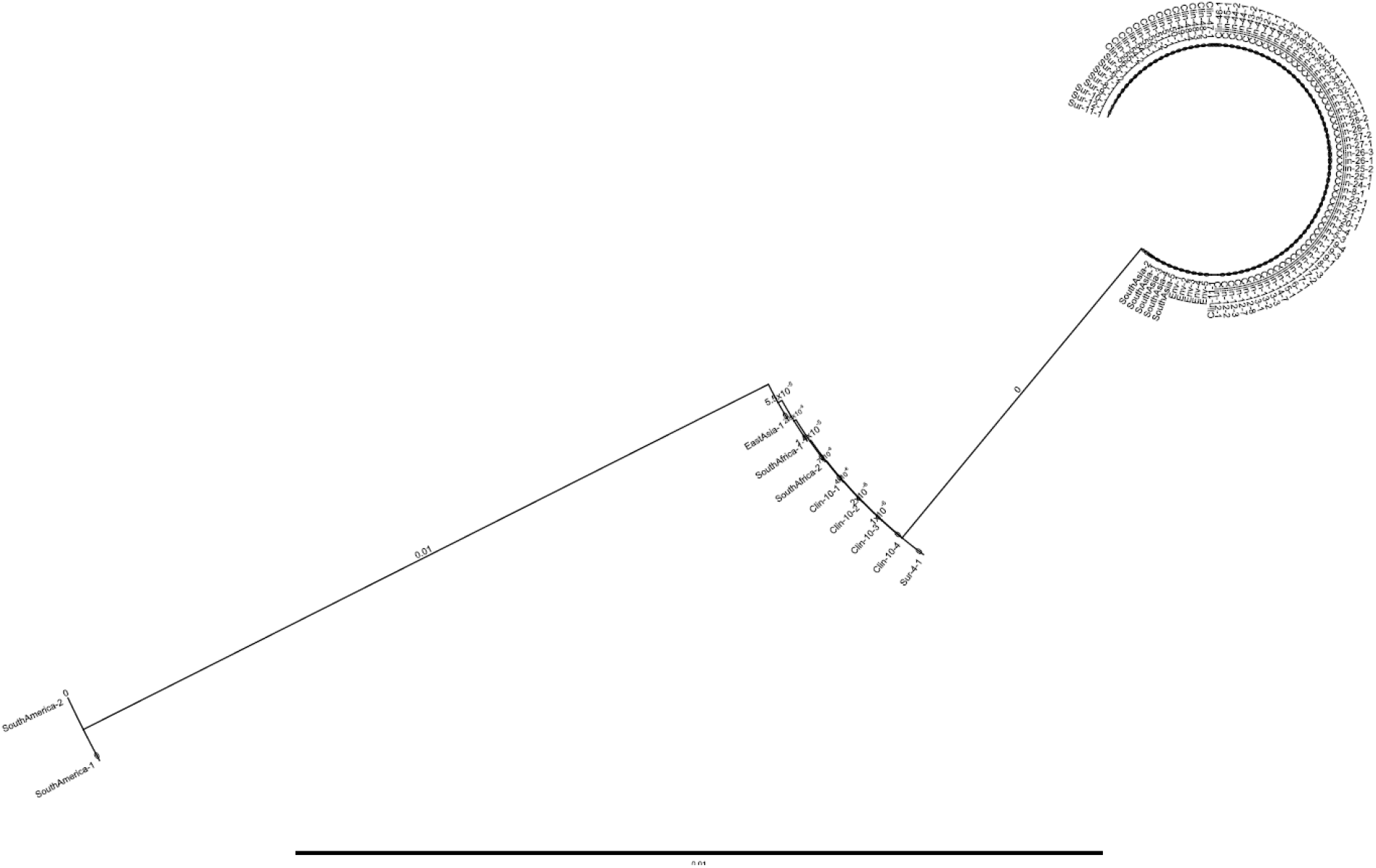
Phylogenetic analysis of *C. auris* isolates from NY using D1/D2 sequences. D1/D2 nucleotide sequences of 622 isolates of *C. auris* recovered from clinical patients (Clin), colonized patients (Sur), environmental surfaces (Env), and 10 standard isolates from the CDC-AR bank were aligned. The neighbor joining method was used to construct the phylogenetic tree. The bootstrap scores are based on 2,000 reiterations. The NY outbreak was predominated by South Asia Clade I with a minor population of East-Asia Clade II. A tree representing a few *C. auris* isolates from each group is shown. Note, D1/D2 could not differentiate South Africa Clade III from East Asia Clade II.

Analysis of 622 clinical, surveillance, and environmental isolates clearly revealed that the outbreak comprised a major genotype of South Asia clade I with a minor genotype of the East Asia clade II in NY. The other two well-known genotypes of *C. auris* including South Africa Clade III and South America Clade IV were not found in NY.

### Antifungal susceptibility testing

A total of 966 *C. auris* isolates were subjected to antifungal susceptibility testing. These included 277 first clinical isolates from 277 clinical patients, 116 subsequent isolates from 74 of those 277 clinical patients, 215 patient surveillance isolates from 48 of the 277 clinical patients, 240 randomly selected patient surveillance isolates from colonized patients, and 95 environmental isolates (Table 5). Of 277 first clinical isolates, one isolate was susceptible to all the antifungals tested while the rest of the 276 isolates were resistant to fluconazole (≥32.0). Of the fluconazole resistant isolates, 224 (81%) had an elevated minimum inhibitory concentration (MIC) to voriconazole (MIC ≥2.0), 170 (61%) were resistant to amphotericin B (MIC ≥2.0), and 2 (0.7 %) were resistant to 5-FC (≥32.0). None of the isolates were resistant to echinocandins.

**Table 5.** Antifungal susceptibility data of *C. auris* isolates recovered from clinical cases, colonized cases and the environment

| **A. First clinical Isolate (#277) from 277 clinical patients** |  |  |  |  |  |  |  |  |  |  |
| --- | --- | --- | --- | --- | --- | --- | --- | --- | --- | --- |
|  | MIC (µg/ml) |  |  |  |  |  |  |  |  |  |
|  | FLU | IZ | VOR | POS | ISA | CAS | MF | AND | AMB | FC |
| Tentative resistance breakpoint | 32.0 | NA | NA | NA | NA | 2.0 | 4.0 | 4.0 | 2.0 | NA |
| MIC<sub>50</sub> | >256 | 0.5 | 2 | 0.25 | 0.5 | 0.12 | 0.12 | 0.25 | 2 | 0.094 |
| MIC range | 8 - ≥256 | 0.12-1 | 0.25-4 | 0.03-1 | 0.05-2 | 0.008- >16.0 | 0.03-0.5 | 0.12 - 0.5 | 0.125-3 | 0.023 - >32 |
| # (%) resistant | 276 (99) | NA | 224 (81) | NA | NA | 0 | 0 | 0 | 170 (61) | 2 (0.7)* |
| **B. Subsequent clinical isolate (#116) from 74 clinical patients** |  |  |  |  |  |  |  |  |  |  |
| MIC<sub>50</sub> | >256 | 0.5 | 2 | 0.25 | 0.5 | 0.12 | 0.12 | 0.25 | 2 | 0.094 |
| MIC range | 128 - ≥256 | 0.12-1 | 0.5-4 | 0.03-0.5 | 0.12-2 | 0.008- >16.0 | 0.03-4 | 0.12 - 0.5 | 0.125-3 | 0.023 - >32 |
| # (%) resistant | 116 (100) | NA | 96 (83) | NA | NA | 3 (2.6) | 4 (3.4) | 3 (2.6) | 67 (58) | 6 (5.1) |
| **C. Surveillance isolates (#215) from 48 clinical patients** |  |  |  |  |  |  |  |  |  |  |
| MIC<sub>50</sub> | ≥256 | 0.5 | 2 | 0.25 | 1 | 0.06 | 0.12 | 0.25 | 2 | 0.064 |
| MIC range | 8 - ≥256 | 0.05 - 1 | 0.125 - 4 | 0.015 - 1 | 0.12 - 2 | 0.008 - 2 | 0.03 - 4 | 0.03 - 4 | 0.19 - 4 | 0.016 - >32 |
| # (%) resistant | 214 (99) | NA | 175 (81) | NA | NA | 6 (2.8) | 6 (2.8) | 6 (2.8) | 143 (67) | 1 (0.5) |
| **D. Surveillance isolates (#240) from colonized patients** |  |  |  |  |  |  |  |  |  |  |
| MIC<sub>50</sub> | >256 | 0.5 | 2 | 0.25 | 0.5 | 0.06 | 0.12 | 0.25 | 2.0 | 0.064 |
| MIC range | 4-≥256 | 0.015-1 | 0.015-8 | 0.008-1 | 0.008-2 | 0.015-0.5 | 0.03-0.5 | 0.03-0.5 | 0.5-3.0 | 0.016->32 |
| # (%) resistant | 239 (99) | NA | 199 (83) | NA | NA | 0 (0) | 0 (0) | 0 (0) | 160 (67) | 5 (2.1) |
| **E. Surveillance isolates (#95) from environment** |  |  |  |  |  |  |  |  |  |  |
| MIC<sub>50</sub> | >256 | 0.5 | 2 | 0.5 | 1 | 0.25 | 0.25 | 0.25 | 2.0 | 0.064 |
| MIC range | 8 - ≥256 | 0.25-1 | 0.06-4 | 0.03-1 | 0.03-4 | 0.015-1 | 0.06-0.5 | 0.08-1 | 0.25-3.0 | 0.032-0.19 |
| # (%) resistant | 94 (99) | NA | NA | NA | NA | 0 (0) | 0 (0) | 0 (0) | 59 (62) | 0 (0) |

MIC = minimum inhibitory concentration; MICs for azoles and echinocandins are defined as the lowest drug concentration that caused 50% growth inhibition compared to the drug-free controls; MICs for amphotericin B and flucytosine are defined as the lowest concentration at which there was 100% and 90% growth inhibition, respectively. MIC<sub>50</sub> was defined as the MIC value at which ≥50% of the isolates of *C. auris* tested were inhibited; FLU = fluconazole, VOR = voriconazole, POS = posaconazole, ISA = isavuconazole, IZ = itraconazole, CAS = caspofungin, MF = micafungin, AND = anidulafungin, AMB = amphotericin B, 5-FC = 5 flucytosine

Of the 116 subsequent clinical isolates analyzed from 74 clinical patients, four (3.4%) were resistant to micafungin (≥4.0); two were recovered from urine, and one each was recovered from blood and ascites from four individual patients. Of two urine isolates with micafungin resistance, one also showed resistance against caspofungin (≥2.0) and anidulafungin (≥4.0). Of echinocandin resistant isolates, two isolates from two patients exhibited borderline resistance to amphotericin B (MIC = 1.5), but CDC’s Mycotic Diseases Branch found these isolates to be susceptible to amphotericin B (≤1.0). Of the follow-up surveillance isolates analyzed from some of the clinical cases, one patient’s 6 isolates were resistant to echinocandins, and of these one isolate recovered from a rectal swab collected to assess for ongoing colonization after resolution of infection was pan-resistant as it also showed resistance to amphotericin B (MIC =3.0), which was retrospectively confirmed in 2019 by CDC’s Mycotic Diseases Branch (MIC =1.5). Interestingly, all urine isolates from this patient collected on different days were resistant to echinocandins, but isolates recovered from other body sites were variably resistant or susceptible (Table 6) indicating the possibility of presence of heterogenous populations. None of the isolates recovered either for colonized patients or from the environment showed any resistance to echinocandins (i.e., echinocandin resistance was only identified in patients meeting the clinical case definition).

**Table 6.**
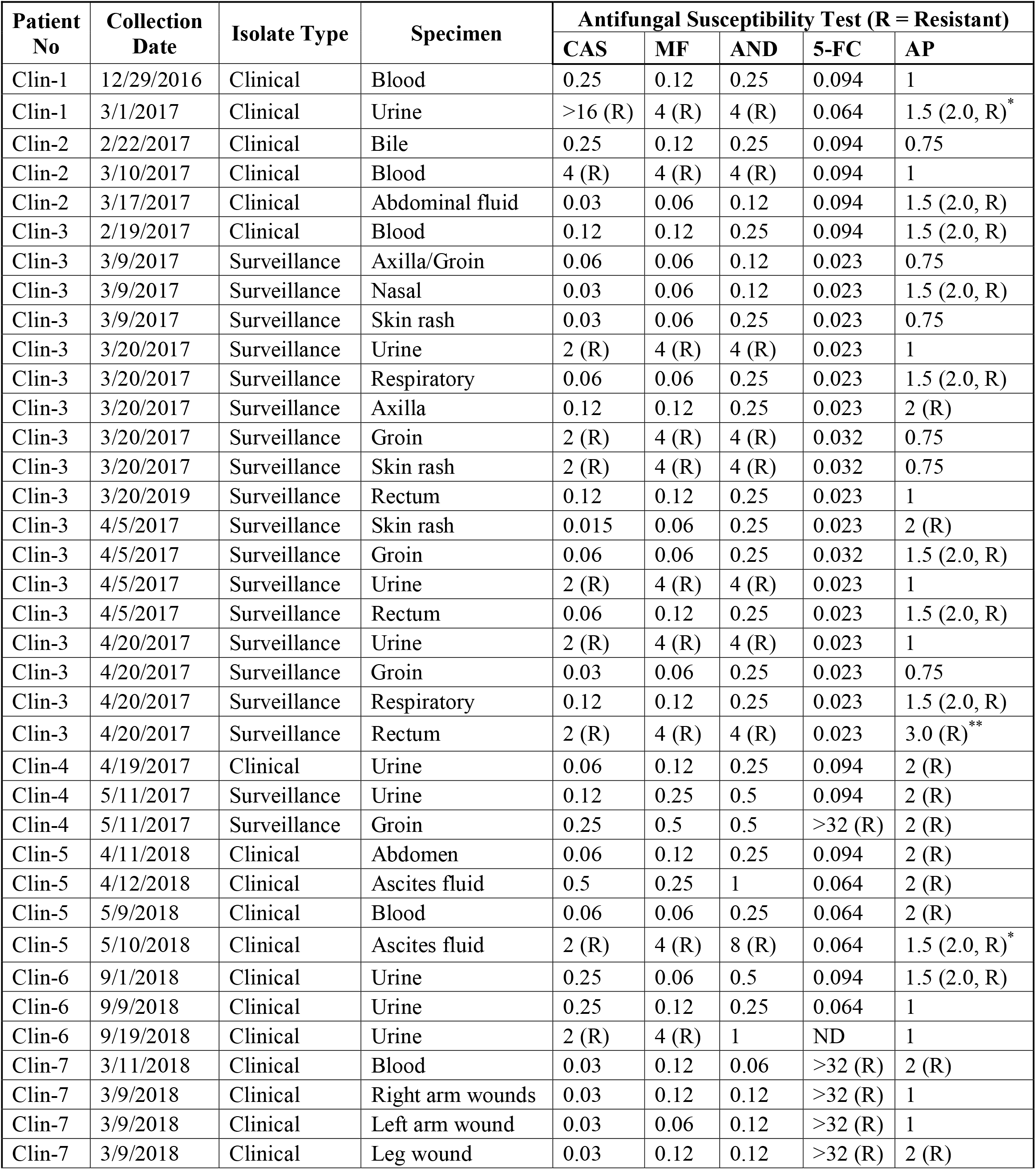
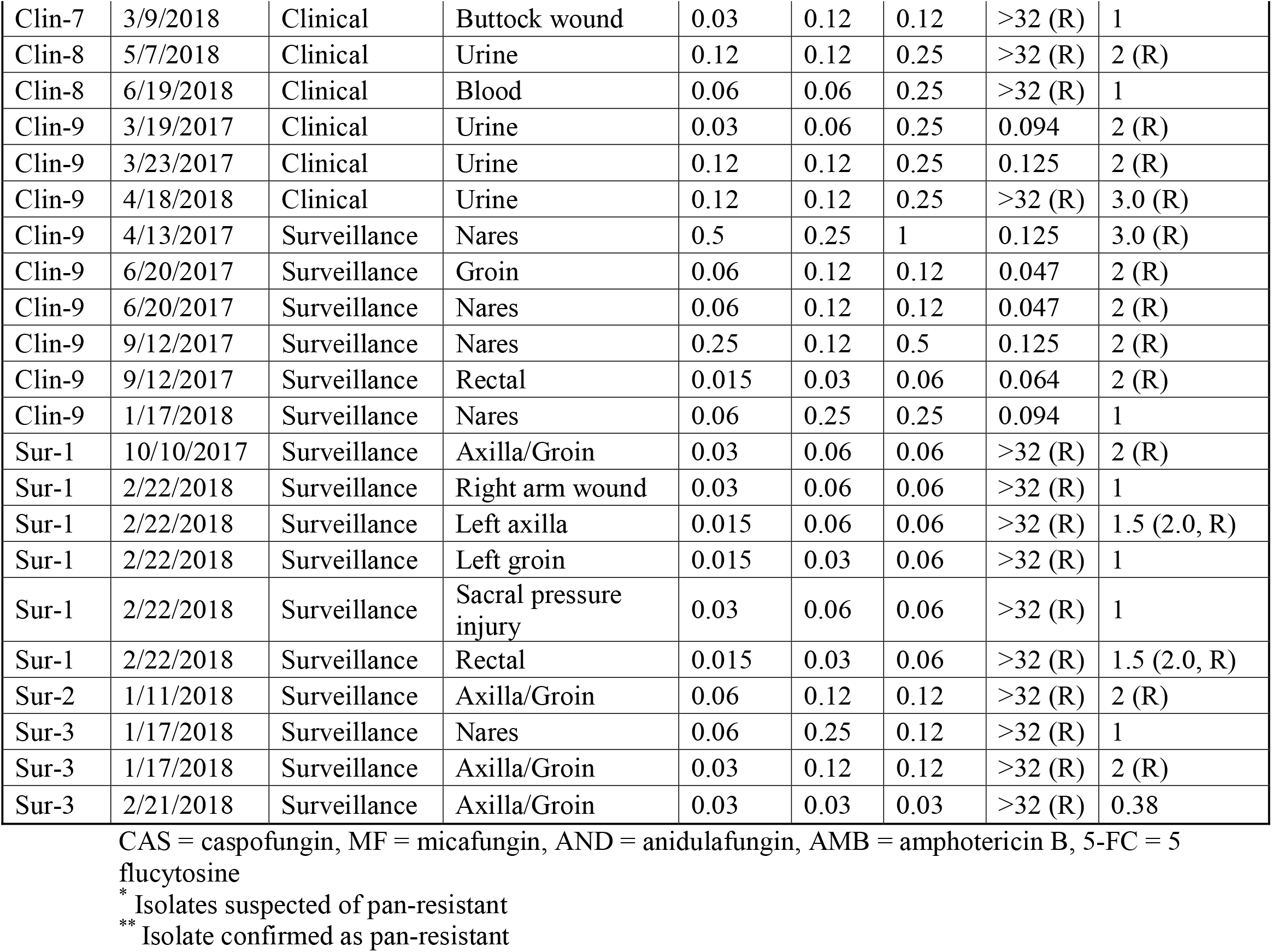
List of serial *C. auris* patient isolates with resistance to various antifungals and/or pan-resistance.

### Isolation of other fungal species from patient and environmental surveillance samples

We determined what other organisms were present in the patient and environmental surveillance samples. *Candida albicans* (13.94%) was by far the dominant pathogen in patient surveillance samples followed by *C. auris* (9.54%), *C. glabrata* (6.49%), *C. tropicalis* (3.98%) and *C. parapsilosis* (3.76%). Several other rare *Candida* spp., other yeasts, molds, and bacterial species were also isolated (Fig. 4 A). The environmental samples were predominantly positive for *C. parapsilosis* (7.03%) followed by *C. auris* (5.42%), *C. albicans* (2.27%), *C. guilliermondii* (2.1%), and *C. glabrata* (1.61%). Other *Candida* species, yeast, molds, and bacteria were also isolated (Fig. 4 B).

**Figure 4 A.**
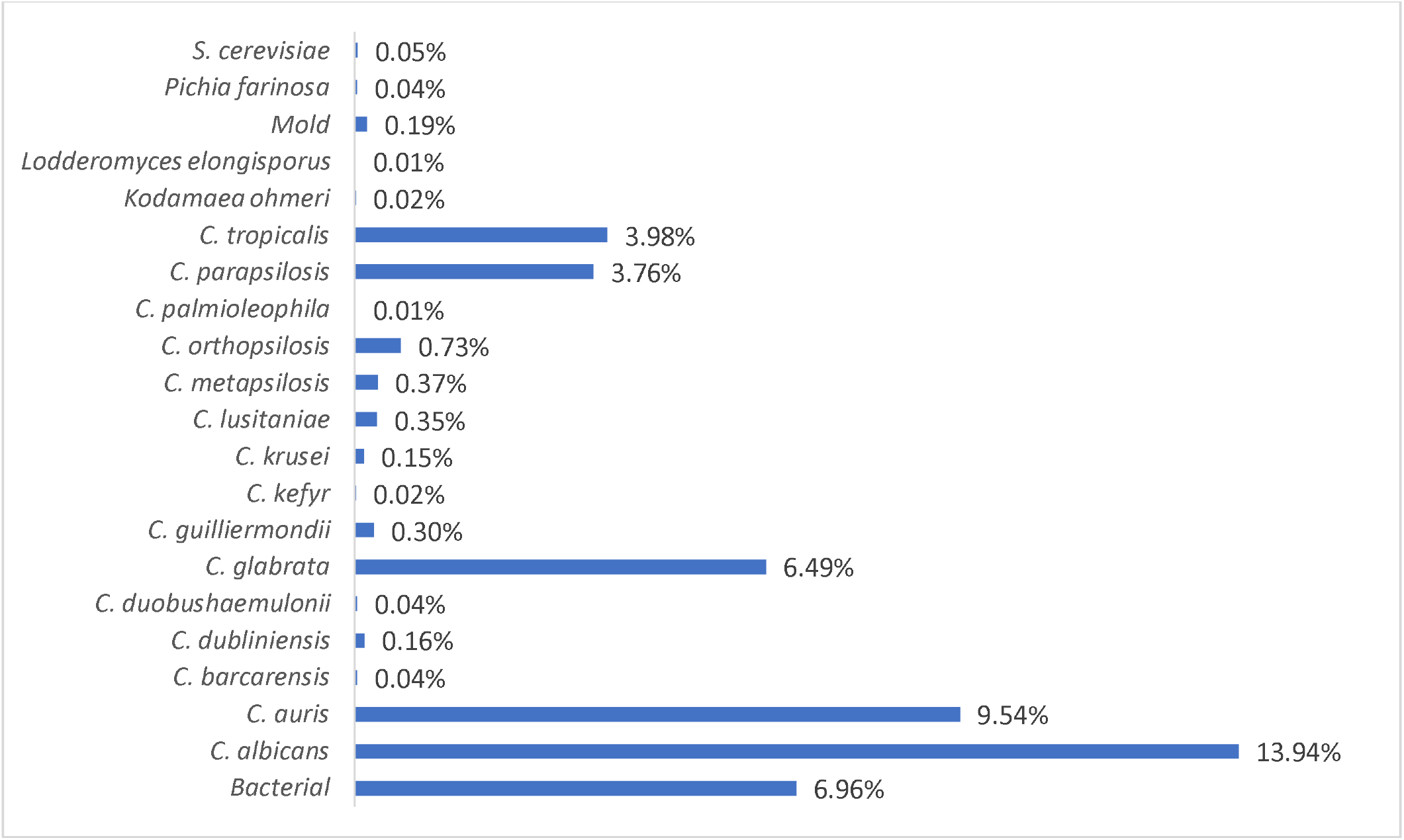
Prevalence of *Candida* species in patient surveillance samples. The bar diagram represents different *Candida* species isolated from patient surveillance samples. *Candida albicans* was the dominant pathogen followed by *C. auris, C. glabrata, C. tropicalis* and *C. parapsilosis*. Other *Candida* species, and other yeasts were minor components.

**Figure 4 B.**
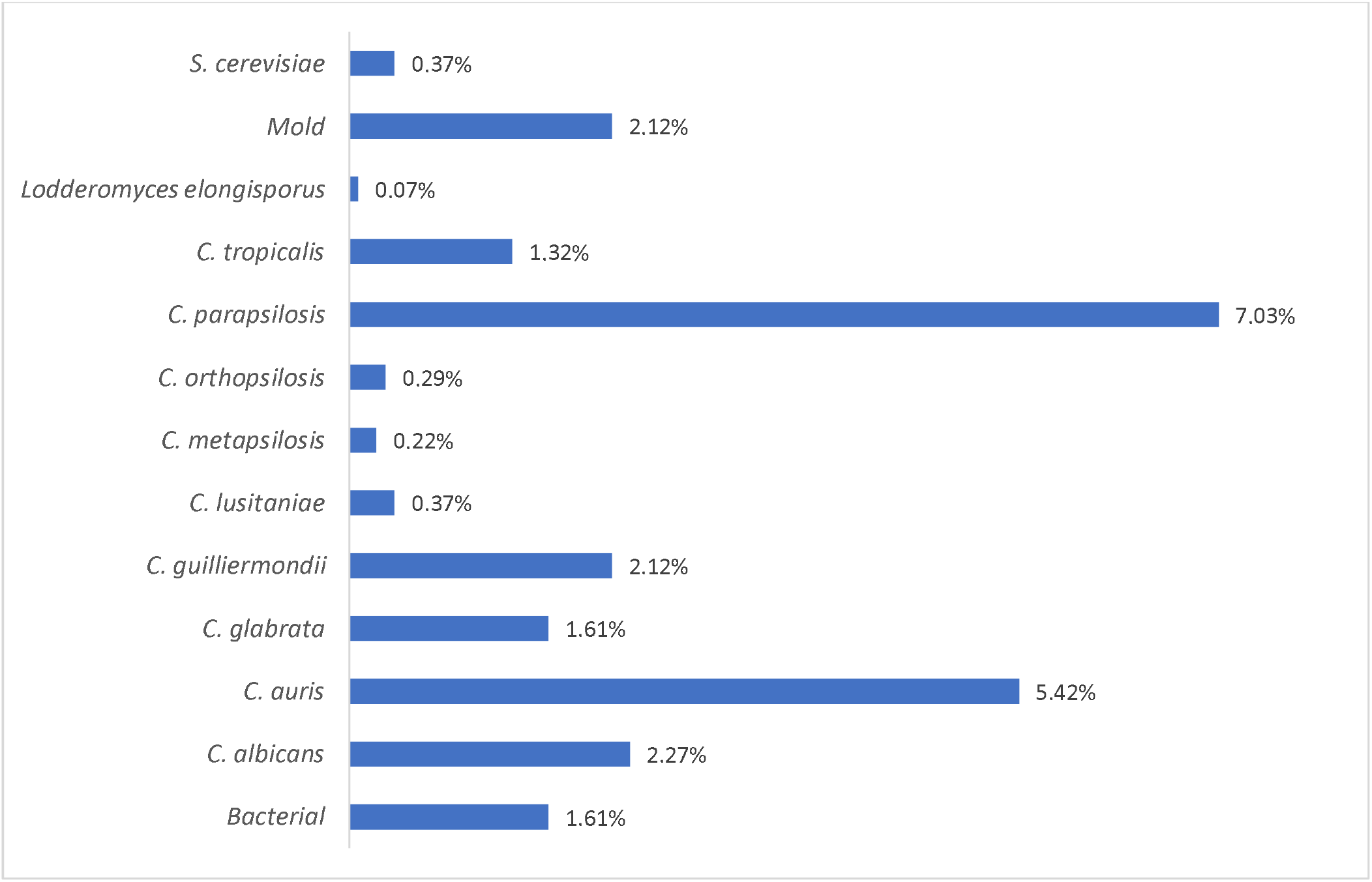
Prevalence of *Candida* species in environmental surveillance samples. Bar diagram represents different *Candida* species isolated from environmental surveillance samples. *Candida parapsilosis* was the dominant pathogen followed by *C. auris, C. albicans, C. guilliermondii, C. glabrata,* and *C. tropicalis.* Other *Candida* species, molds, and other yeasts were minor components.

## DISCUSSION

*Candida* species are major causes of healthcare-associated infections in the United States (22). Infections by these pathogens result in mortality rates of >30-40% and are responsible for the highest total annual hospitalization costs of any invasive fungal disease (n = 26,735; total cost $1.4 billion) (22–24). The recent emergence of multidrug-resistant (MDR) *C. auris* has further exacerbated this crisis (9, 16). Since 2016, New York State and New York City metropolitan areas have borne the brunt of this unprecedented epidemic. Despite coordinated efforts of public health authorities, *C. auris* infections continue to increase across New York State. We report the largest outbreak of *C. auris* in New York with 277 clinical and 350 colonized cases identified from August 2016 through 2018. As of July 18, 2019, the outbreak has impacted 173 healthcare facilities in which the patients received care in the 90 days prior to their *C. auris* diagnosis were affected. These including 64 hospitals, 106 nursing homes, 1 LTACH and 2 hospices.

Detecting outbreaks of infectious diseases at an early stage is crucial for timely implementation of infection control measures and for minimizing morbidity and mortality. The real-time PCR assay developed recently by our laboratory (21) proved to be an excellent diagnostic tool for the rapid detection of *C. auris* from patient surveillance samples with high sensitivity and accuracy. As compared to patient surveillance samples, the calculated diagnostic accuracy and specificity of the real-time PCR assay for environmental samples was lower. However, these calculations rely on culture as the gold standard, but this is not an appropriate comparator for environmental samples. For example, we did not know how long *C. auris* was present in the healthcare environment prior to sample collection. Also, several environmental samples received for testing were often collected after the site had been cleaned and disinfected. The ability of real-time PCR assay to pick up DNA from live, dead and growth defective *C. auris* as well as its ability to pick up leftover DNA stuck to the objects likely impacted the calculated diagnostic accuracy. Nevertheless, environmental testing contributed substantially to the institution of infection control practices in the affected healthcare facilities. We would recommend that while PCR is an excellent screening tool for testing of environmental samples, culture should be used as the basis for follow up remediation efforts.

Despite extensive efforts, the outbreak is ongoing. This could be due to the complexity of the healthcare systems in the city including density and close proximity of healthcare facilities in the region, and/or the lack of testing availability within facilities. The Wadsworth Center Mycology Laboratory has taken on the burden of essentially all testing since the beginning of the outbreak, and the volume by itself is a limiting factor in the assessment of an outbreak of this magnitude. It is imperative that more clinical, public, and commercial laboratories adopt *C. auris* testing technologies. Additionally, there is an urgent need for the development of high throughput assays as well as point-of-care molecular tests for the rapid turnaround time of lab testing.

The phylogenetic data using ITS and D1/D2 regions of the ribosomal genes revealed that the South Asia clade I was the major genotype and the East Asia clade II was a minor genotype of the *C. auris* outbreak in NY. These results are consistent with whole genome sequencing results reported for NY strains (25). The distinct separation of South and East Asia clades by ribosomal gene sequencing in the present study is encouraging as next generation sequencing technologies are not in widespread use by clinical, public, and commercial laboratories for fungal pathogens. Interestingly, we did not find any isolate of the South Africa Clade III or the South America Clade IV in NY despite being a large international port of entry. The current method of phylogenetic analysis with ribosomal genes is well established for the separation of South Asia Clade I, East Asia Clade II, and South America Clade III with the exception of South Africa Clade IV. Using ITS sequences, South Africa Clade IV clustered with South Asia Clade I consistent with an earlier study (5) but it clustered with East Asia Clade I using D1/D2 sequences. In summary, ribosomal genes served as an excellent marker for the separation of all well-known clades of *C. auris*.

In the present laboratory-based aggregate analysis, we quantified *C. auris* in colonized patients and their surrounding environment to determine the extent of colonization. Our results revealed that the colonized patients harbored large numbers of live *C. auris* on skin (axilla/groin) and mucosal (nares) surfaces. Likewise, the patient’s contact points including several healthcare objects were also contaminated with the large numbers of live *C. auris*. These results indicate that patients who are heavily colonized with *C. auris* on their skin or mucosal surfaces can contaminate their surroundings, which might be a key to the successful transmission of *C. auris* in the healthcare facilities. Identifying reservoirs, prompt notification of *C. auris* identification, and implementation of effective infection control practices are key to effectively controlling the spread of *C. auris*.

Among colonized patients, the nares appeared to serve as an excellent site for *C. auris* growth as it harbored an approximately 2-logs higher CFU than the axilla/groin. The precise mechanism(s) leading to the extensive colonization of nares is currently unclear, but results of this investigation led us to use one combination swab of nares/axilla/groin rather than a separate individual swab for colonization screening, an approach necessary to handle an outbreak of this magnitude. Further research is needed to understand the mechanism(s) of colonization. Although, *C. auris* recovery from the patient’s skin and healthcare environment have been established previously (8, 9, 16), in this report, we have used a quantitative approach to determine the extent of *C. auris* colonization of skin and mucosal surfaces of the colonized patients and their surrounding environment.

The resistance to fluconazole, voriconazole, and amphotericin B observed in the current outbreak was higher than the two large studies previously published (15, 20) and this discrepancy could be due to the fact that *C. auris* outbreak in NY was dominated by South Asia Clade I. South Asia Clade I was intrinsically resistant to fluconazole and approximately 81% of these isolates had elevated MIC to voriconazole and 61% to amphotericin B. The echinocandin resistance was noted in fewer isolates, which was consistent with other studies (15, 20). Pan-resistance is always a concern in *C. auris,* and despite the large outbreak, we found only three isolates suspected to have pan-resistance based on testing done at the Wadsworth Center Mycology Laboratory with one isolate ultimately confirmed as pan-resistance at the CDC’s Mycotic Diseases Branch. Our MIC values for two suspected pan-resistant isolates that did not confirm were 1.5 while CDC’s Mycotic Diseases Branch reported them to be lower than 1.5. There are more data emerging that indicate that amphotericin B resistance is inducible and transient, and MIC values of some isolates decrease following passage in the laboratory (26). Amphotericin B resistance is rare in most of the *Candida* species and it is thought to be associated with a fitness cost (27). In contrast, closely related species to *C. auris*, *C. haemulonii*, *C. duobushaemulonii* and *C. pseudohaemulonii* have high-level intrinsic resistance to amphotericin B (20) and therefore possibly a contributing factor to rare infections worldwide. The borderline resistance observed in *C. auris* against amphotericin B is intriguing and suggests it has the ability to overcome fitness costs in order to survive, spread and cause infection worldwide. More research is needed to understand the mechanisms leading to amphotericin B resistance in *C. auris*. In our investigation, we have found a small percentage of isolates showing resistance to 5-flucytosine, which is indicative of either the patient having received combination therapy of amphotericin B and 5-flucytosine or the spread of resistant isolates in the healthcare facilities. Nevertheless, fewer isolates showing echinocandin or 5-FC resistance among the many isolates tested in the current outbreak clearly indicate that these isolates did not spread extensively in the healthcare facilities. More studies are needed to understand the impact of resistance on the fitness and survival of the isolates.

In summary, use of a real-time PCR assay as a rapid detection tool, quantification of the extent of colonization of *C. auris* on biotic (patient) and abiotic (environment) surfaces, successful use of one combination swab of nares/axilla/groin to combat the large outbreak of *C. auris* in New York, and the use of ribosomal genes to determine relatedness among *C. auris* isolates were among some of the highlights of this investigation and provide further insight into the *C. auris* outbreak in NY.

---

**Supplementary Table 1.** Identification of *Candida auris* and other closely related species from yeast isolates submitted by clinical, public and commercial laboratories

| Wadsworth ID<br>(MALDI-TOF MS) | Submitter's ID | Number of<br>Isolates |
| --- | --- | --- |
| *Candida auris* | *C. auris* | 183 |
|  | *C. haemulonii* | 103 |
|  | R/O *C. auris* | 70 |
|  | No. ID | 24 |
|  | Yeast | 24 |
|  | *C. famata* | 1 |
|  | *C. glabrata* | 1 |
|  | *C. guilliermondii* | 1 |
|  | *C. lusitaniae* | 1 |
|  | *Candida* spp. | 5 |
| **Total = 413** |  |  |
| *C. duobushaemulonii* | *C. auris* | 3 |
|  | *C. duobushaemulonii* | 2 |
|  | *C. haemulonii* | 3 |
|  | *C. haemulonii* /*C. duobushaemulonii* | 2 |
|  | Not provided | 1 |
|  | Yeast | 1 |
| **Total = 12** |  |  |
| *C. haemulonii* | *C. haemulonii* | 2 |
|  | Not provided | 1 |
|  | R/O *C. auris* | 3 |
|  | Presumptive *C. auris* | 1 |
| **Total = 7** |  |  |

## ACKNOWLEDGMENTS

We are grateful to members of the Healthcare Epidemiology and Infection Control Program and staff members from various hospitals and long-term-care facilities for assistance with surveillance samples. We thank Wadsworth Center Clinical Laboratory Reference System for funding and Wadsworth Center Applied Genomic Technologies and Media & Tissue Cultures Cores for DNA sequencing and culture media, respectively. We are also thankful to Kimberly McClive-Reed for editorial comments. This publication was supported by Cooperative Agreement number NU50CK000516, funded by the Centers for Disease Control and Prevention. Its contents are solely the responsibility of the authors and do not necessarily represent the official views of the Centers for Disease Control and Prevention or the Department of Health and Human Services.

## Authors Contributions

Yan Zhu was responsible for the processing of the surveillance sample, culture identification of all yeasts and data analyses. Lynn Leach was responsible for the real-time PCR assay of patient surveillance samples and data analysis; Brittany O’Brien performed antifungal susceptibility testing of all *C. auris* isolates; Alex Clarke and Mariam Bates were responsible for the analysis of environmental samples using real-time PCR assay; Deborah Blog, Eleanor Adams, Belinda Ostrowsky, Karen Southwick, Richard Erazo, Elizabeth Dufort, Valarie B. Haley, Coralie Bucher, and Monica Quinn assisted in epidemiology investigation of positive results, including advising and arranging for onsite point prevalence surveys, performing onsite infection control assessments and contact tracing. Eleanor Adam, Belinda Ostrowsky, and Monica Quinn also critically read the manuscript. Ron Limberger coordinated the lab-epi meeting and critically read the manuscript. Emily Lutterloh coordinated the epidemiologic *C. auris* response and critically reviewed the manuscript. Vishnu Chaturvedi provided his expertise in antifungal susceptibility testing and critically reviewed the manuscript. Sudha Chaturvedi conceived and designed the study, interpreted data, and wrote the manuscript.

## References

1. Satoh K, Makimura K, Hasumi Y, Nishiyama Y, Uchida K, and Yamaguchi H. *Candida auris* sp. nov., a novel ascomycetous yeast isolated from the external ear canal of an inpatient in a Japanese hospital. Microbiol Immunol. 2009;53(1):41–4.

2. Lee WG, Shin JH, Uh Y, Kang MG, Kim SH, Park KH, et al. First three reported cases of nosocomial fungemia caused by *Candida auris*. Journal of clinical microbiology. 2011;49(9):3139–42.

3. Chowdhary A, Sharma C, Duggal S, Agarwal K, Prakash A, Singh PK, et al. New clonal strain of *Candida auris*, Delhi, India. Emerging infectious diseases. 2013;19(10):1670–3.

4. Wang X, Bing J, Zheng Q, Zhang F, Liu J, Yue H, et al. The first isolate of *Candida auris* in China: clinical and biological aspects. Emerging microbes & infections. 2018;7(1):93.

5. Magobo RE, Corcoran C, Seetharam S, and Govender NP. *Candida auris*-associated candidemia, South Africa. Emerging infectious diseases. 2014;20(7):1250–1.

6. Emara M, Ahmad S, Khan Z, Joseph L, Al-Obaid I, Purohit P, et al. *Candida auris* candidemia in Kuwait. Emer Infect Dis. 2014;21(6):1091–2.

7. Calvo B, Melo AS, Perozo-Mena A, Hernandez M, Francisco EC, Hagen F, et al. First report of *Candida auris* in America: Clinical and microbiological aspects of 18 episodes of candidemia. The Journal of infection. 2016;73(4):369–74.

8. Schelenz S, Hagen F, Rhodes JL, Abdolrasouli A, Chowdhary A, Hall A, et al. First hospital outbreak of the globally emerging *Candida auris* in a European hospital. Antimicrobial resistance and infection control. 2016;5:35.

9. Vallabhaneni S, Kallen A, Tsay S, Chow N, Welsh R, Kerins J, et al. Investigation of the First Seven Reported Cases of *Candida auris*, a Globally Emerging Invasive, Multidrug-Resistant Fungus - United States, May 2013-August 2016. MMWR Morbidity and mortality weekly report. 2016;65(44):1234–7.

10. Ben-Ami R, Berman J, Novikov A, Bash E, Shachor-Meyouhas Y, Zakin S, et al. Multidrug-Resistant *Candida haemulonii* and *C. auris,* Tel Aviv, Israel. Emerg Infect Dis. 2017;23(1):195–203.

11. Morales-Lopez SE, Parra-Giraldo CM, Ceballos-Garzon A, Martinez HP, Rodriguez GJ, Alvarez-Moreno CA, et al. Invasive Infections with Multidrug-Resistant Yeast *Candida auris*, Colombia. Emerging infectious diseases. 2017;23(1):162–4.

12. Ruiz Gaitan AC, Moret A, Lopez Hontangas JL, Molina JM, Aleixandre Lopez AI, Cabezas AH, et al. Nosocomial fungemia by *Candida auris*: First four reported cases in continental Europe. Revista iberoamericana de micologia. 2017;34(1):23–7.

13. Spivak ES, and Hanson KE. *Candida auris*: an Emerging Fungal Pathogen. Journal of clinical microbiology. 2018;56(2).

14. Chowdhary A, Anil Kumar V, Sharma C, Prakash A, Agarwal K, Babu R, et al. Multidrug-resistant endemic clonal strain of *Candida auris* in India. European journal of clinical microbiology & infectious diseases: official publication of the European Society of Clinical Microbiology. 2014;33(6):919–26.

15. Lockhart SR, Etienne KA, Vallabhaneni S, Farooqi J, Chowdhary A, Govender NP, et al. Simultaneous Emergence of Multidrug-Resistant *Candida auris* on 3 Continents Confirmed by Whole-Genome Sequencing and Epidemiological Analyses. Clinical infectious diseases: an official publication of the Infectious Diseases Society of America. 2017;64(2):134–40.

16. Adams E, Quinn M, Tsay S, Poirot E, Chaturvedi S, Southwick K, et al. *Candida auris* in Healthcare Facilities, New York, USA, 2013-2017. Emerging infectious diseases. 2018;24(10):1816–24.

17. Chowdhary A, Sharma C, and Meis JF. *Candida auris*: A rapidly emerging cause of hospital-acquired multidrug-resistant fungal infections globally. PLoS Pathog. 2017;13(5):e1006290.

18. Jeffery-Smith A, Taori SK, Schelenz S, Jeffery K, Johnson EM, Borman A, et al. *Candida auris*: a Review of the Literature. Clinical microbiology reviews. 2018;31(1).

19. Mizusawa M, Miller H, Green R, Lee R, Durante M, Perkins R, et al. Can Multidrug-Resistant *Candida auris* Be Reliably Identified in Clinical Microbiology Laboratories? Journal of clinical microbiology. 2017;55(2):638–40.

20. Kathuria S, Singh PK, Sharma C, Prakash A, Masih A, Kumar A, et al. Multidrug-Resistant Candida auris Misidentified as Candida haemulonii: Characterization by Matrix-Assisted Laser Desorption Ionization-Time of Flight Mass Spectrometry and DNA Sequencing and Its Antifungal Susceptibility Profile Variability by Vitek 2, CLSI Broth Microdilution, and Etest Method. Journal of clinical microbiology. 2015;53(6):1823–30.

21. Leach L, Zhu Y, and Chaturvedi S. Development and Validation of a Real-Time PCR Assay for Rapid Detection of Candida auris from Surveillance Samples. Journal of clinical microbiology. 2018;56(2).

22. Strollo S, Lionakis MS, Adjemian J, Steiner CA, and Prevots DR. Epidemiology of Hospitalizations Associated with Invasive Candidiasis, United States, 2002-2012(1). Emerging infectious diseases. 2016;23(1):7–13.

23. Magill SS, O’Leary E, Janelle SJ, Thompson DL, Dumyati G, Nadle J, et al. Changes in Prevalence of Health Care-Associated Infections in U.S. Hospitals. The New England journal of medicine. 2018;379(18):1732–44.

24. Benedict K, Jackson BR, Chiller T, and Beer KD. Estimation of direct healthcare costs of fungal diseases in the United States. Clinical infectious diseases: an official publication of the Infectious Diseases Society of America. 2018.

25. Chow NA, Gade L, Tsay SV, Forsberg K, Greenko JA, Southwick KL, et al. Multiple introductions and subsequent transmission of multidrug-resistant *Candida auris* in the USA: a molecular epidemiological survey. The Lancet Infectious diseases. 2018;18(12):1377–84.

26. Lockhar SR. Candida auris and multidrug resistance: Defining the new normal. Fungal Genetics & Biology. 2019;131.

27. Vincent BM, Lancaster AK, Scherz-Shouval R, Whitesell L, and Lindquist S. Fitness trade-offs restrict the evolution of resistance to amphotericin B. PLoS biology. 2013;11(10):e1001692.

